# 4-Dimensional Chess: Acoustic Localisation Reveals Nested Spatio-temporal Strategies in an Arboreal Communication Network

**DOI:** 10.64898/2026.09.21.752596

**Authors:** Edmund W. Basham, Luke C. Larter, Patrick S. Champagne, Douglas L. Jones, Timothy H. Keitt

**Author notes:** Denotes joint first authorship (Edmund W. Basham, Luke C. Larter).

## Abstract

1. Adaptive behavioural strategies require animals to simultaneously navigate social and ecological domains across multiple spatial and temporal scales. Although drones and computer vision have recently transformed the study of wild animal societies, many nocturnal species and those occupying structurally complex habitats remain inaccessible to these approaches, limiting our understanding of behaviour in natural settings.
2. We aimed to determine how behavioural strategies are organised across nested spatial and temporal scales within a wild communication network.
3. We used three-dimensional acoustic localisation and source separation to track individual male *Hyperolius sp. A*, a nocturnal African reed frog, within a natural rainforest chorus and quantify patterns of site fidelity, movement, spatial organisation, and call-timing interactions.
4. Males exhibited significant site fidelity across nights, while chorus spatial structure varied with local caller density. Within nights, individuals followed a stereotyped behavioural sequence, descending from elevated arboreal refugia before settling into lower calling positions near breeding sites. At finer temporal scales, call-timing interactions varied according to both local competitor density and the proximity of neighbouring rivals.
5. These findings demonstrate that behavioural strategies emerge across nested spatial and temporal scales and that long-term spatial positioning, short-term movement decisions, and moment-to-moment signalling interactions are tightly linked within natural communication networks. More broadly, acoustic localisation provides a powerful framework for studying behaviour in species and habitats that remain difficult to observe using conventional approaches.

## 1. INTRODUCTION

Adaptive social strategies require animals to navigate complex social environments by coordinating various behaviours with conspecifics in space and time (Alexander, 1974). Furthermore, ecological challenges such as acquiring resources, avoiding predators, and tracking appropriate habitat, require complex spatiotemporal navigation of an animal’s environment (Morse, 1980). Both social and ecological processes are inextricably linked, with a species’ ecology influencing the frequency, location, and tenor of interactions with conspecifics (Krebs & Davies, 1978). This means that individuals must navigate both domains simultaneously, and must do so across several nested temporal and spatial scales (Webber et al., 2023). However, laboratory experiments are infeasible at the spatiotemporal scales at which most wild animals live their lives, and the difficulty of collecting data in the field often requires trade-offs between how detailed observations are and the scales over which data can be collected. Thus, developing a holistic understanding of how adaptive strategies play out in the wild across various intertwined spatiotemporal scales remains a key challenge in behavioural ecology.

Anuran choruses are a perfect test case for how broader strategies unfold across nested spatiotemporal scales. Breeding occurs in aggregations that are typically highly localized in both space and time, with choruses forming at bodies of water just after nightfall throughout a well-defined annual breeding season (Gerhardt & Huber, 2002; Wells, 1977). Male strategies then play out around breeding sites over a variety of temporal scales. In species with prolonged breeding seasons, chorus attendance at the seasonal scale is a key determinant of annual reproductive success, and this arises as the summation of numerous nightly chorus-attendance decisions (Friedl & Klump, 2005). At the scale of a single night, males must navigate to breeding sites from their daytime refuges at the appropriate time and, when calling sites are not permanent, must claim and defend productive calling sites from rivals (Wells, 1977). Thus, the spatial structure of the chorus is typically dynamic within and across nights, as it emerges from ongoing negotiations among rivals as they partition calling habitat, defend territories, and maintain and enforce inter-male spacing patterns (Dougherty et al., 2022; Wilczynski & Brenowitz, 1988; Wells, 1978; Brenowitz, 1989).

Finally, at the most fine-grained temporal scales, male success depends on successfully outcompeting rivals while calling to attract females. Males compete to produce calls with the most attractive acoustic properties, and to call at attractive times relative to the calls of rivals (Dyson et al., 2013; Greenfield, 1994; Larter and Ryan, 2025). Furthermore, vocal competition and spatial negotiations are interwoven strands of a more expansive communication strategy. Males defend calling sites to reduce local acoustic interference and competition, while also primarily interacting vocally with nearer neighbours who pose the greatest local reproductive threat (Brenowitz, 1989; Brush and Narins, 1989; Schwartz, 1993; Wells, 1977). Thus, anuran choruses have become paradigmatic examples of communication networks (Grafe, 2005; Snijders and Naguib, 2017; Reichert et al., 2021); complex networks of interconnected signallers and receivers in which emergent communication dynamics and a species’ social organization in space and time reciprocally influence one another.

Thus, overall, we see in anurans that successful male social strategies require skilful navigation of space and time at numerous nested and inter-connected scales. Furthermore, in arboreal anurans that call from above-ground vegetation, these social dynamics unfold within a fundamentally three-dimensional environment (Hödl, 1977). Vertical position influences signal propagation, exposure to temperature and humidity gradients, foraging success, and vulnerability to predators, creating trade-offs that may shape calling strategies and site selection (Hödl, 1977; Richards & Wiley, 1980). However, tracking behaviour across multiple spatiotemporal scales in 3D environments presents challenges to traditional field-based methods. Most field studies rely on visual surveys, focal recordings, or short observational windows that cannot continuously track and record vocalisations from individuals in complex 3D habitats (Basham et al., 2022). Consequently, fundamental questions—such as whether individuals maintain consistent spatial positions, how they move within choruses over time, and how spatial relationships shape real-time interactions—remain difficult to address in the wild.

Recent advances in bioacoustic analysis offer a means to overcome these limitations. Distributed microphone arrays and beamformer algorithms exploit differences in signal arrival times of different sound sources across sensors to unmix complex acoustic scenes into separate audio streams per source (Calsbeek et al., 2022; Jones & Ratnam, 2009; Jones et al., 2014, Rhinehart et al., 2020). These approaches can resolve the 3D positions of multiple animals calling simultaneously and assign each call to the individual callers that produced them with high spatial and temporal resolution, even in structurally complex habitats. They therefore provide continuous, non-invasive measurements of both where and when individuals signal, enabling the reconstruction of individual call sequences, the dynamics of calling interactions, and movement trajectories within natural choruses. This creates the opportunity to link spatial positioning, movement dynamics, and signalling interactions within a single empirical framework.

Here, we apply three-dimensional acoustic localisation to wild choruses of an undescribed species of *Hyperolius* from Central African rainforest systems (hereafter *Hyperolius sp. A*), an arboreal reed frog from lowland Central African rainforests that forms dense nocturnal breeding choruses. By tracking changes in 3D chorus structure across nights, individual movement trajectories within nights, and fine-scale moment-to-moment chorusing interaction patterns, we paint a uniquely holistic picture of complex spatio-temporal strategies unfolding in a wild communication network. Our results provide a powerful proof-of-concept that novel bioacoustic technologies can reveal detailed dynamics of previously inaccessible wild animal societies.

## METHODS

### 2.1 Study system and array

Fieldwork was conducted from August to December 2024 at Baposso Village, Ngounié Province, southern Gabon (2.0839° S, 12.13416° E; 650–800 m.a.s.l.; Fig. 1a). The study site comprised dense moist tropical forest surrounding a small vernal pond (Fig. 1b). The pond was surrounded and overhung on all sides by dense understory vegetation of shrubs, deadwood, lianas, and small trees, and was overlooked by a number of larger diameter canopy trees of ∼20 meters in height whose crowns partially intersected above the pond.

**Fig. 1:**
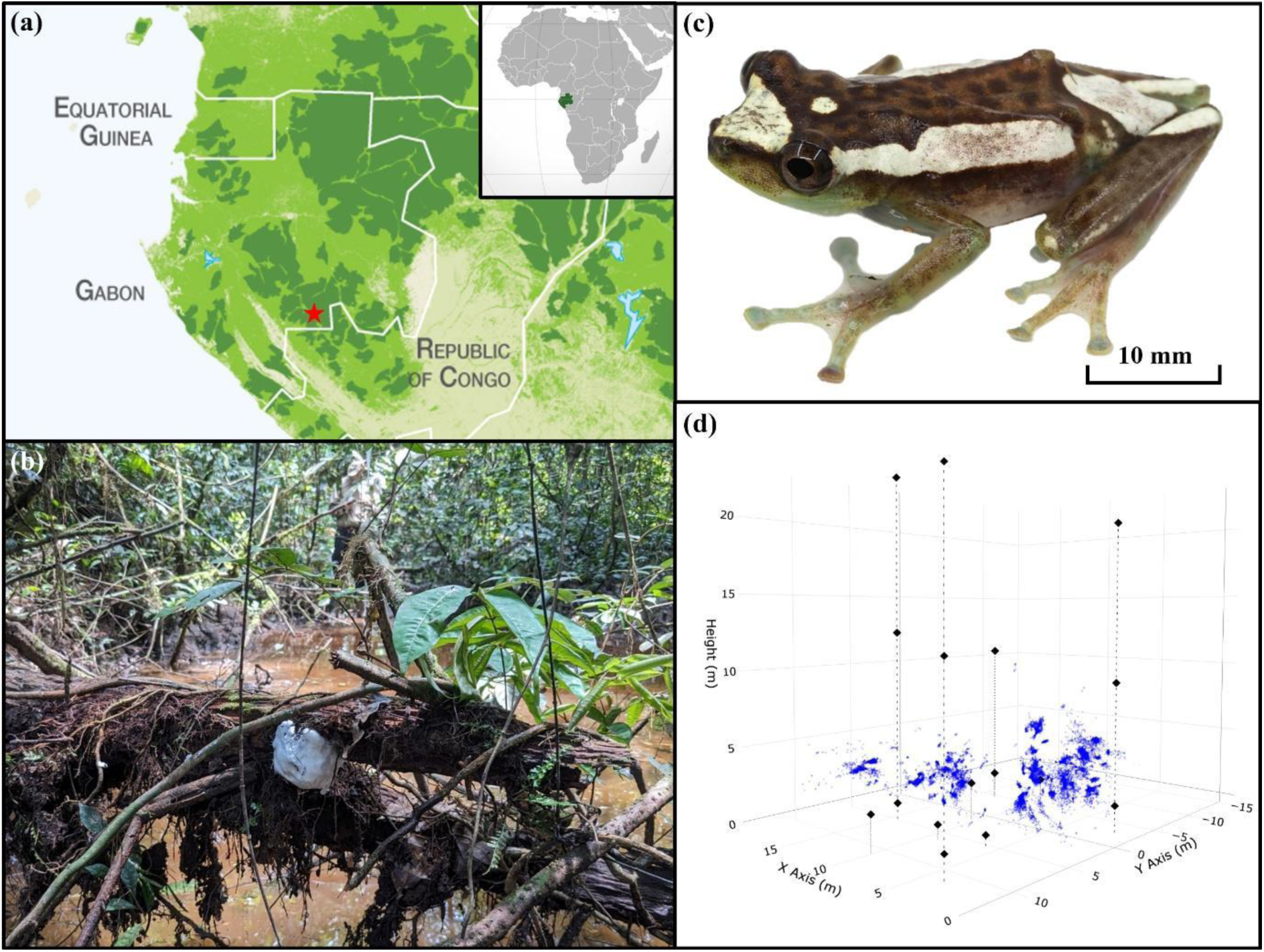
(a) Map of Central African forest cover, showing intact forest landscapes (dark green) and other forest cover (light green). The study site in Gabon is indicated by a red star. Map adapted from GRID-Arendal (https://www.grida.no). (b) The vernal pond where the array was installed. (c) *H. sp. A* (Individual deposition number: NBM-AR-015773). (d) Representation of the 3D array using the *plotly* package (Sievert et al., 2015). Microphones are black diamonds and all triangulated locations of calls are blue points. An interactive version of this plot can be viewed via this link (https://geckoeb.github.io/Frogs3D-Plot/frog_3D_plot.html) or by running the R code provided via the figshare repository.

A three-dimensional microphone array of 14 GPS-integrated Audiomoth devices was deployed around the pond. The array was arranged in a rhomboid configuration, with four vertical stacks extending from the understory (∼2 m) to the canopy (∼20 m), and additional units placed within the understory to capture calling activity near the pond (Fig. 1d). Devices recorded at 16 kHz with variable schedules reflecting field logistics, equipment performance, and calling activity. Analyses focus on two sessions (Session 3: October and Session 4: November 2024) during which *H. sp. A* (Fig. 1c) exhibited peak calling activity as the wet season rains began at the start of October and array performance allowed reliable localisation. All recordings began at 18:00 but durations differed between sessions due to site access logistics which impacted battery replacement, with October recordings lasting 3.5 hours (12,600 s) and November recordings 45 minutes (2,700 s).

Microphone positions were estimated from pairwise distance measurements between all devices using a laser distance meter (Leica Disto D2). The relative positions of microphones in the array were then mathematically reconstructed using these measurements.

### 2.2 Acoustic localisation

Acoustic localisation was performed using a multi-source localisation framework introduced in detail elsewhere (Jones et al., 2014; Jones & Ratnam, 2009), which integrates time-of-arrival estimation, triangulation, clustering, and adaptive beamforming, to resolve the positions and signals of multiple simultaneously calling individuals in complex acoustic environments. The approach is robust to irregular microphone geometry, uncertainty in sensor placement, environmental noise, and overlapping calls, making it suitable for dense natural choruses.

The method first bandpass filters recordings to isolate the focal species and segments the multi-channel signal into short overlapping windows. Within each segment, relative time delays between microphones are estimated and used to compute a least-squares source location for each acoustic event. Segments dominated by a single source, such as when a frog produces a call, are identified using triangulation error and inter-microphone correlation, forming spatial clusters corresponding to that individual caller. A recent addition groups segments by comparing their full sets of microphone-pair time delays, using agglomerative clustering based on Hamming distance, to link segments from the same location. This improves source identification by constraining cluster structure, with final source locations estimated as the mean position of clustered segments.

For each localisation cluster, we defined a robust centre as the median x, y, and z coordinates, and calculated the Euclidean distance of each point from this centre. Points exceeding a threshold of the median distance plus two median absolute deviations (MAD) were excluded. We additionally retained only the closest 50% of points ranked by distance to the centre, ensuring that analyses were based on the densest and most reliable subset of localisations. Clusters with fewer than four points were not filtered.

### 2.3. Statistical analysis

#### (i) Chorus spatial structure

To examine broader-scale movement, 2-minute windows (runs) were sampled at ∼15-minute intervals across recording sessions. Although individual identity could not be continuously verified across these runs, spatial clusters remained sufficiently distinct and moved in logical continuous paths over time, making it highly certain that we were tracking movement of the same individuals through time. Analyses were conducted in R (v4.3.1) (R Core Team, 2023).

For spatial analysis of the chorus, we computed the average 3D location of each male within a night; i.e. the average x, y, and z coordinates corresponding to all calls produced and located. Within-night movement was therefore treated as positional uncertainty around a stable calling site. We also calculated the centroid of the x, y, and z coordinates for each male in each run, to allow us to quantify the precision of our triangulation procedure. Within a run, we calculated the Euclidean distance between each individual triangulation point and this centroid and then calculated the standard deviation (SD) of these distances as a measure of radial error. We then calculated the mean radial error across all runs for all males in all nights to provide an overall measure of 3D precision.

To quantify the spatial structure of the chorus, we calculated the Euclidean distance between the night-centroids of all males calling together at the chorus. To test whether spatial organization varied with chorus density, we fitted two linear models at the night level, with chorus size (number of callers detected per night) as the predictor variable and either mean nearest-neighbour distance or mean calling height as the response variable. Each night therefore contributed a single observation to each model. Then, to assess site fidelity without individual identification, we tested whether calling locations were more spatially consistent across nights than expected under a null model. Spatial consistency between nights was quantified by optimally matching nightly sets of frog centroids in three-dimensional space using the Hungarian algorithm (Kuhn, 1955), and calculating the mean distance between matched locations. When chorus sizes differed between nights, unmatched individuals were excluded. Null expectations were generated by random sampling from the empirically occupied calling space (Supplementary Methods 1).

Observed distances were compared to a permutation null (999 iterations) in which centroids were randomly sampled from the voxel space. Empirical p-values were calculated as the proportion of null distances less than or equal to the observed value. To assess temporal decay in site fidelity, we extended this analysis across all within-session night pairs (lags 1–10 days), testing whether mean matched distances increased with time.

#### (ii) Modelling within-night vertical movement

Visual inspection of individual trajectories suggested a stereotyped within-night pattern, in which males descend from elevated refugia early in a session before settling into more stable, lower calling positions for its remainder. We tested this descent-and-settle hypothesis in two stages: first, by characterising the discrete behavioural states underlying vertical repositioning; and second, by directly testing whether vertical movement declined over the course of a session (see below). To characterise these behavioural states, Hidden Markov Models (HMMs) were fitted to the vertical height (z, metres) time series of 109 individual frog × night combinations (frog-nights). Session duration varied between sampling periods (45 min or 3.5 h), such that longer sessions contributed more position fixes and therefore more behavioural transitions for HMM fitting. All position fixes were calculated while males were actively calling; HMM states therefore describe the magnitude of vertical repositioning between consecutive calls. Our approach relies on acoustic localization and so does not allow us to track movements while animals are not actively vocalizing. A 3-state model was selected on the basis of statistical fit and biological plausibility (see Table S1). All analyses were conducted using the *moveHMM* package (Michelot et al., 2016).

The HMM was fitted to step lengths using a gamma distribution with no angular component, as movement direction was not recorded. Initial parameter values are provided in Table S2. State decoding was performed using the Viterbi algorithm via the fathom and stateProbs functions in *moveHMM*, with no constraints placed on transition probabilities during fitting. Three behavioural states were identified: Active Displacement (large step lengths indicative of energetic canopy transit); Local Repositioning (low-to-moderate step lengths representing fine-scale repositioning); and Stationary (minimal positional change consistent with settled calling). Model selection between two- and three-state HMMs used AIC; the three-state model was strongly preferred (ΔAIC = 27.96; Table S1), consistent with guidance that state number should reflect ecological plausibility (Pohle et al., 2017).

Prior to statistical modelling, two sequential filters were applied to the behavioural state dataset. Frog-nights with fewer than 5 total observations were excluded (n = 14 removed), and frog-nights shorter than 1,800 s (30 minutes) were excluded (n = 4 removed). Importantly, HMM state assignment was performed on the complete unfiltered dataset; filters were applied exclusively at the modelling stage. The final dataset comprised 1,051 observations from 91 frog-nights across 20 recording sessions.

Three models were employed to interrogate complementary aspects of the descent-and-settle hypothesis. A linear model (LM) was fitted with signed vertical displacement (dz, m) as the response and session elapsed time (minutes) as the predictor, testing whether net vertical movement is significantly negative early in sessions. Two binomial GLMMs were fitted with Active Displacement (yes/no) and Stationary (yes/no) as respective responses, each with session elapsed time (hours) as the fixed-effect predictor, testing whether active displacement declines and stationary calling increases across the night. Time was expressed in hours for GLMMs to avoid numerical scaling issues. The two GLMMs included random intercepts for frog-night (1 | ID). The linear model was fitted using base R; the two binomial GLMMs were fitted using *lme4* (Bates et al., 2015) and *lmerTest* (Kuznetsova et al., 2017). Additional details on the HMM pipeline are provided in Supplementary Methods 2.

#### (iii) Chorusing Interactions

To characterise fine-scale interactions, we analysed six continuous recording periods ranging from 12.5 to 20 minutes in length from within the first 60 minutes of each of six nights (the 10th, 11th, and 12th of October, and the 5th, 6th, and 7th of November; Fig. S1). Periods were chosen when *H. sp. A* chorusing intensity was high and background noise was relatively low (Fig. S1).

The predominant call type of *H. sp. A* is a short (mean duration 33 ± 7 ms) tonal call lacking frequency modulation (mean carrier frequency 2.13 ± 0.13 kHz) (Fig. 2a). These ‘single-note calls’ are the most prevalent call type, with >96% of calls in the current study being single-note calls. However, males also occasionally produce ‘multi-note calls’ (Fig. 2a).

**Fig. 2:**
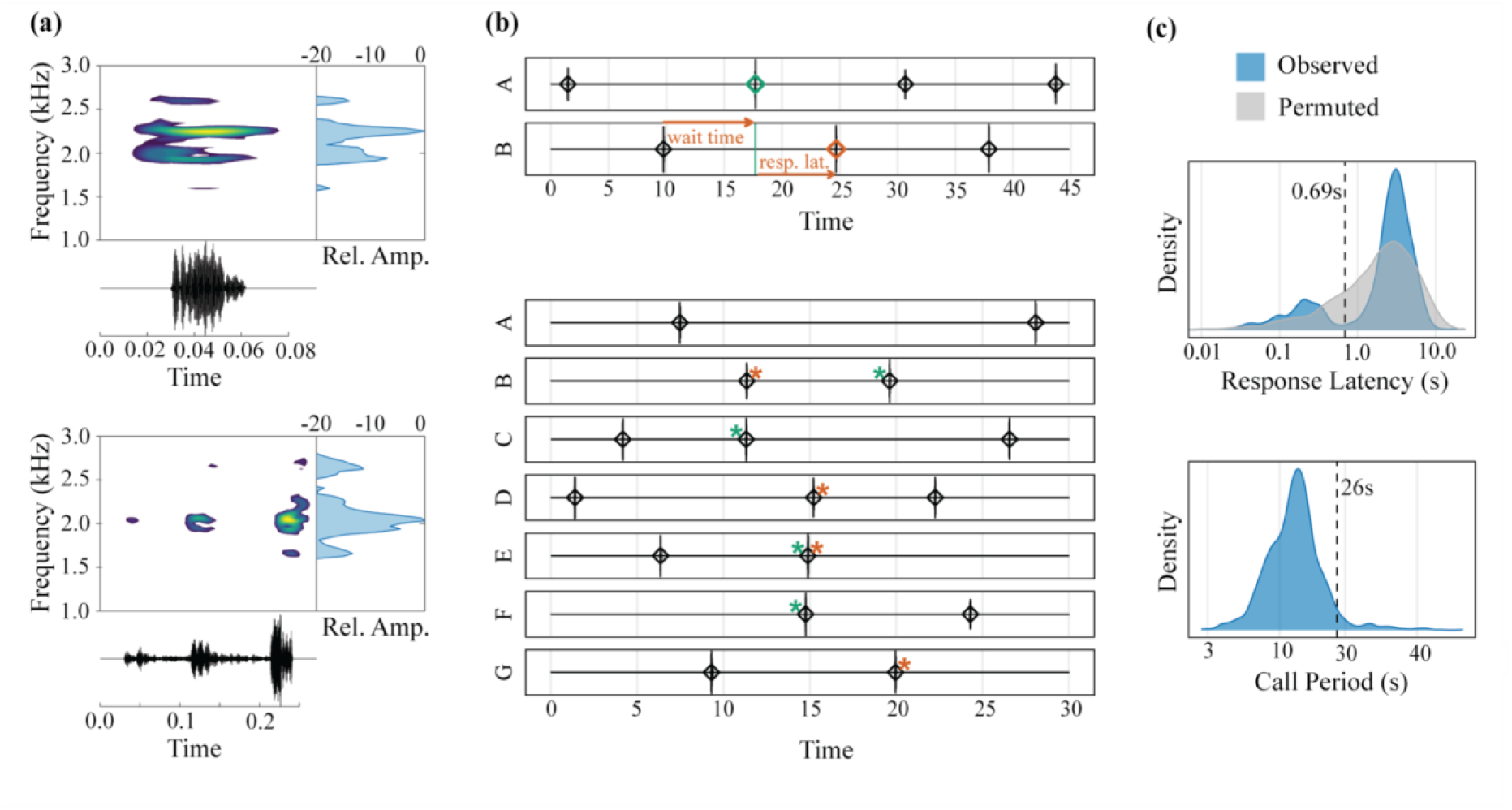
(**a**) *Upper*: Spectrogram, waveform, and power spectra (Relative Amplitude) of a single-note call. *Lower*: multi-note call. Spectrograms were generated with the spectro() function from the *SeeWave* R package (window length = 512, overlap = 80%). Brighter colours denote higher relative amplitudes. (**b**) *Upper*: Visualization of 45s of calling interactions between 2 males. The thin vertical black lines with black diamonds represent the calls. A transition in the chorus sequence in which one male (orange) responds to the call of another (green) is highlighted to illustrate important temporal variables; ‘wait time’ is the interval between the responding male’s most recent previous call and the rival’s call to which he is now responding (green), and ‘response latency’ is the interval between this rival’s call (green) and the responding male’s response to it (orange). *Lower*: Visualization of 30s of a 7-male chorus. Short-latency ‘rapid responses’ are denoted with asterisks; green asterisks to the left denote the preceding male of the interaction, and orange asterisks the responding male. Back-to-back rapid responses are visible at 15s. (**c**) *Upper*: Density plot of log-transformed response latencies pooled across all 6 recording nights. Blue distribution is observed response latencies, grey is the permutation derived null. The line at 0.69s denotes the threshold used to differentiate rapid and slow responses, based on k-means clustering. *Lower*: Density plot of log-transformed call periods, pooled across recordings. The line at 26s denotes the 95% percentile of call periods.

##### Analysis of Chorusing Dynamics

Chorusing dynamics were analysed across nights spanning a range of caller densities. The onsets of calls were difficult to identify due to background noise present in our recordings, so we used the times of peak amplitude of calls as their timestamps. These peaks were readily identifiable, and the brevity of this species’ calls (∼33 ms) means peak times are a good proxy for call onset times. For multi-note calls, we used the time of peak amplitude of the first call note.

Here, a male calling in response to a rival’s call could respond with one of two discrete response types. Responding males could respond with lengthy call latencies (*call latency*: the time elapsing between that rival’s call and the responding male’s response call), or respond with rapid, sub-second call latencies (Fig. 2b). This generated a response latency distribution with clear bimodality, which was absent in a permutation-derived null distribution (Fig. 2c). This demonstrates that these two clusters represent discrete active response types, rather than incidental social by-products. Cluster analysis revealed that 0.69 s was the optimal threshold for separating these two response types (Supplementary Methods 3). For subsequent analysis, we categorized responses as ‘rapid’ or ‘slow’ based on this threshold.

##### Modelling the Drivers of Male Call-Timing Behaviour

We used mixed effects models (*lme4*: (Bates et al., 2015)) to model how endogenous rhythms and social factors influenced calling responses. We modelled call sequences within choruses as successive transitions from one caller to the next unfolding through time. For each observed transition from one caller to the next, we then asked; given that male B called directly after male A in the call sequence, what influenced whether they did so with a rapid response or a slow response? And, within each response type, what influenced B’s response latency? This transition-by-transition approach has proven successful in studying the complex temporal dynamics of frog choruses (Larter & Ryan, 2024b). Not all males called consistently during recordings. Thus, we excluded calls from analysis when they were preceded by exceptionally long call periods (*call period*: the time elapsing between successive calls by the same caller). This was because these lengthy intervals likely represent males ceasing to call for a time, rather than call periods generated while actively calling. We used 26s as our threshold, as this was the 95% percentile of our log-transformed call period distribution, and visually approximated the upper limit of the bulk of call periods (Fig. 2c). We also use this same threshold when generating a time-varying measure of chorus activity levels (*number of active callers*, below).

To analyse the probability that males responded to a rival’s call with a rapid or slow response, we built a mixed effects logistic regression with response type (rapid or slow) as the response variable (n = 2,193) and included several time-varying fixed effects. *Wait time*: the time that had elapsed between the responding male’s most recent prior call and the rival’s call to which he was currently responding (to control for cyclical changes in responsiveness between calls; (Larter & Ryan, 2024b)). *Number of active callers*: the number of unique rivals that had called within the 26 seconds prior to the current response call (to control for differing levels of chorus activity within and between nights). *Inter-caller distance*: the Euclidean distance in meters between the responding male and the rival he was responding to. We did not include call type (single vs. multi-note) of response calls or responded-to rival’s calls as a predictor variable as multi-note calls were rare (75 out of 2256 calls = 3.6%) and biased towards a few very active individuals.

Regarding random effects, we could not definitively identify whether individuals appearing on different recording nights were the same individuals or not. Thus, we treated each frog-night combination as a unique ID, and included random intercepts for responding male ID and responded-to male ID nested within recording date. We acknowledge that this could lead to repeated measures of the same individuals being treated as distinct subjects on different nights. However, we consider this a more conservative approach than relying on potentially erroneous cross-night identity assignments, which could introduce serious bias. We also included all random slopes supported by the data.

We then ran separate Gaussian models to investigate whether the latency of responses within each response type was influenced by these same fixed effects (slow response model: n = 1694; rapid response model: n = 469). For slow responses, we log-transformed response latencies to improve model fit. To investigate whether males increase their call rates in response to local social dynamics, we built a Gaussian model with call period as the response variable (n = 2193) and the same fixed effects structure as above. We also included whether each call was a rapid or slow response, to reveal how response type influenced the resulting call period. Finally, to investigate whether call periods were more variable in more active choruses, we constructed a Gaussian model with the coefficient of variation of call period as the response variable. We calculated the coefficient of variation (CV) of call periods for each male for each number of active callers’ value he experienced and regressed this on the number of active callers (the number of unique rivals that had called within the last 26 seconds, defined above). Here, we only included CVs calculated from at least 10 calls (n = 63).

## RESULTS

### 3.1 Chorus Spatial Structure

Across the 20 recording nights, between 4 and 8 individual *H. sp. A* were detected per night (mean ± SD: 5.4 ± 1.2 frogs per night; n = 109 frog-nights in total). Mean nearest-neighbour distance between simultaneously calling individuals was 5.07 ± 1.87 m (range: 1.75 – 11.1 m; Fig. S2a) and mean calling height across all frog-nights was 3.33 ± 1.44 m (range: 1.34 – 10.4 m; Fig. S2b). Across 1,118 frog-run-nights, the mean SD of radial distances from the run centroid was 0.0765 m, indicating high repeatability of localisation estimates within runs.

Mean nearest-neighbour distance declined significantly with increasing chorus size (linear regression: β = −0.608 m per additional frog, t₁₈ = −3.77, R² = 0.44, p = 0.001; Fig 3.), indicating that frogs were packed more closely together. Mean calling height showed a positive trend with chorus size, though this relationship fell just short of conventional significance (β = +0.180 m per additional frog, t₁₈ = 2.04, R² = 0.19, p = 0.056). This suggests a weak tendency for frogs to call from greater heights on nights with larger choruses, though the evidence is marginal and should be interpreted cautiously.

**Fig. 3:**
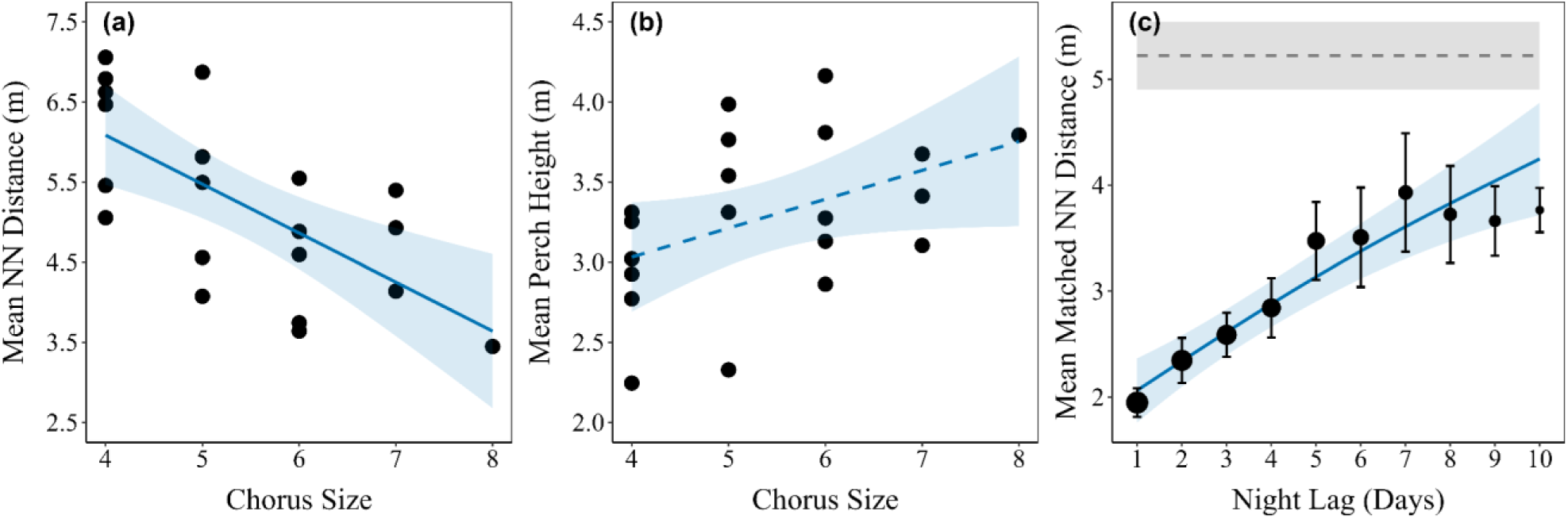
Spatial structure of the *H. sp. A* calling chorus across 20 recording nights. (a) Mean nearest-neighbour distance as a function of chorus size; solid regression line indicates a significant result. (b) Mean calling height as a function of chorus size; dashed regression line indicates marginal result. (c) Temporal decay of site fidelity: mean distance between optimally matched frog-night centroids as a function of night lag (≤10 days). Points show observed values (size ∝ number of night pairs, from 18 at lag = 1 to 3 at lag = 10), with ±1 SE error bars. The solid blue line shows the fitted GAMM trajectory. The grey ribbon and dashed line indicate the null expectation (±2 SD and mean, respectively) from 999 permutations. Shaded blue bands in (a, b, and c) show 95% confidence intervals.

Frogs exhibited strong spatial consistency across nights, with calling locations far more similar than expected under random habitat use (mean observed=1.97 m vs mean permuted = 5.24 m; p < 0.001). However, this spatial consistency declined with increasing time between nights. Across within-session comparisons (1–10 day lags), mean matched distances increased significantly with night lag (GAMM, p < 0.001), indicating a gradual decay in site fidelity over time.

### 3.2 HMM State Classification

A total of 1,125 vocalisation events were recorded across 109 frog-nights. Step sizes ranged from 0.0004 m to 5.17 m (mean 0.35 m). Following sequential filtering, 1,051 observations from 91 frog-nights were retained for statistical modelling.

The three-state HMM resolved a clear gradient in vertical repositioning between consecutive vocalisation events. State 1 (Stationary) represented near-fixed calling positions (mean step 0.054 ± 0.048 m), State 2 (Local Repositioning) moderate positional adjustments (0.339 ± 0.334 m), and State 3 (Active Displacement) large-scale vertical movement consistent with canopy transit (0.870 ± 0.932 m). Local Repositioning was the dominant state (56.2%, n = 632), followed by Stationary (24.8%, n = 279) and Active Displacement (19%, n = 214). Mean calling height differed across states, with Active Displacement highest (3.55 ± 1.68 m), followed by Stationary (3.43 ± 1.20 m) and Local Repositioning (2.98 ± 1.33 m; Table S3). Transition probabilities are provided in Table S4; Fig. S4.

Temporal dynamics revealed consistent within-night progression in behavioural state composition (Fig. 4a). Active Displacement was predominant at session onset, declining rapidly to near absence within the first hour of recording. Local Repositioning dominated throughout the remainder of the session, consistently accounting for most observations. Stationary increased progressively across the session, rising from a low proportion at session onset to representing a minority of observations by mid-to-late session. These shifts coincided with a decline in mean calling elevation from 3.8 m to 2.5–3.0 m after which elevation stabilised. Signed elevation change was predominantly negative in early observations before converging on zero.

**Fig. 4.**
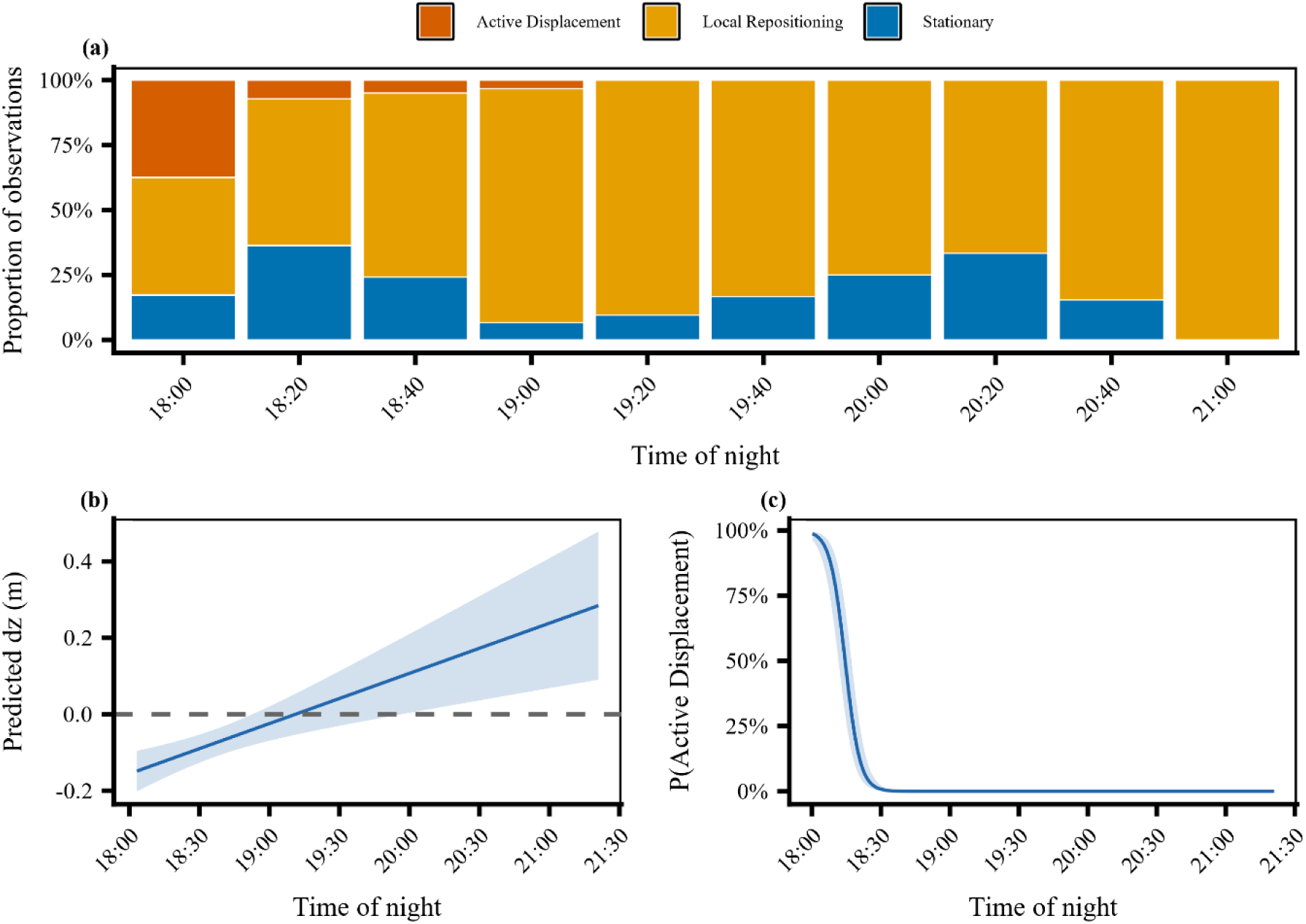
a) Behavioural state composition of male *H. sp. A* across nightly recording sessions (18:00–21:30). Bars represent the proportion of observations assigned to each of three HMM-decoded behavioural states (Active Displacement, orange; Local Repositioning, gold; Stationary, blue) within 20-minute time bins. (b) Predicted signed elevation change (dz, m) from the LMM. The solid line represents the population-level mean and the shaded region the 95% confidence interval; the dashed horizontal line marks zero. (c) Predicted probability of Active Displacement from the binomial GLMM.

A linear model confirmed an initial net downward movement at session onset (intercept = −0.156 m, SE = 0.028, t = −5.507, p < 0.001), followed by a return toward zero displacement over time (β = +0.00219 m min⁻¹, SE = 0.0006, t = 3.706, p < 0.001) (Fig. 4b). Generalised linear mixed models showed that the probability of Active Displacement was initially high (intercept = +4.456 log-odds, SE = 0.812, z = 5.489, p < 0.001) but declined rapidly within 30 minutes (OR = 9.10 × 10⁻⁹, z = −8.009, p < 0.001) (Fig. 4c). In contrast, Stationary increased over time, from 6% at session onset (intercept = −2.777 log-odds, SE = 0.399, z = −6.966, p < 0.001) to 35% by 3.5 hours, with odds increasing by a factor of 1.92 per hour (z = 2.678, p = 0.007) (Fig. S5). The three modelling approaches produced complementary results, collectively supporting a structured temporal progression in behavioural state across the session.

### 3.3 Chorusing Behaviour

#### (i) Drivers of Response Latency

Longer wait times (GLMM: β = 0.46 ± 0.06 SE, z = 7.83, p < 0.001; Fig. 5a), and greater numbers of actively calling rivals (GLMM: β = 0.61 ± 0.1 SE, LRT, z = 6.19, p < 0.001; Fig. 5b) significantly increased the probability that males responded with rapid responses. Furthermore, responses to nearer rivals were significantly more likely to be rapid responses (GLMM: β = -0.81 ± 0.12 SE, z = -6.79, p < 0.001; Fig. 5c). Longer wait times led to significantly faster response latencies within rapid responses (LMM: β = -0.02 ± 0.006 SE, t = -3.3, p = 0.001) and slow responses (LMM: β = -0.07 ± 0.009 SE, t = -7.69, p < 0.001).

**Fig. 5:**
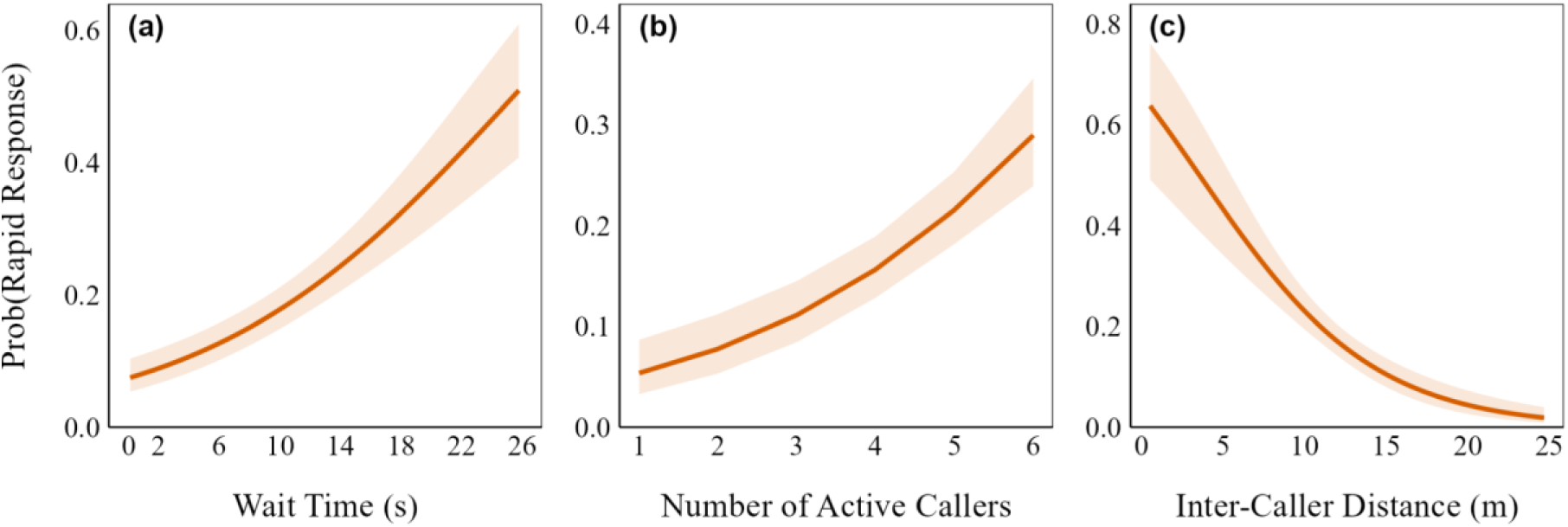
Estimated marginal mean effects of different predictor variables on the probability that males responded to a rival’s call with a rapid response (calling < 0.69s after that rival’s call) vs. a slow response (≥ 0.69s).

Greater numbers of actively calling rivals led to significantly faster response latencies within slow responses (LMM: β = -0.24 ± 0.027 SE, t = -8.67, p < 0.001), but not within rapid responses (LMM: β = 0.009 ± 0.007 SE, t = 1.24, p = 0.21). Unexpectedly, slow responses were significantly slower when responding to nearer rivals (LMM: β = -0.1 ± 0.018 SE, t = -5.41, p < 0.001), while rapid responses showed a non-significant trend towards being faster when responding to nearer rivals (LMM: β = 0.016 ± 0.008 SE, t = 1.93, p = 0.07). Estimated marginal mean effects of predictor variables for all models relating to call timing are presented in Supplemental Table S5.

#### (ii) Call Periods

Choruses were consistently active throughout the recordings with no apparent structuring of calling interactions into bouts, leading to a unimodal distribution of log-transformed call periods (Fig. 2c). Individual males would occasionally drop out of the chorus for a time, but not in a coordinated way.

Call periods that ended in rapid responses were significantly shorter than call periods that ended in slow responses (LMM: β = -0.96 ± 0.24 SE, t = -4.08, p < 0.001), but only by ∼1 second on average (call period resulting from slow response = 13.24 s; rapid response = 12.27 s; see Table 1). The mean call period overall was 12.8 s [95% CI: 11.7, 13.8], thus this only represents a reduction of ∼8% of a typical call period. When controlling for response type, a greater number of actively calling rivals did not significantly influence call period duration (LMM: β = -0.32 ± 0.21 SE, t = -1.5, p = 0.124). However, call periods that ended in responses to nearer rivals showed a slight and non-significant trend towards being shorter (LMM: β = 0.23 ± 0.12 SE, t =1.87, p = 0.062; Table 1). The mean coefficient of variation in call period was 0.33 [95% CI: 0.3, 0.35], and there was a slight and nonsignificant trend towards males exhibiting higher call period CVs when a greater number of rivals were actively calling (LMM: β = 0.02 ± 0.01 SE, t = 1.87, p = 0.061; estimated marginal means [95% CI] of call period CV when different numbers of rivals active: 2 rivals, 0.29 [0.25, 0.34]; 4 rivals, 0.32 [0.29, 0.35]; 6 rivals, 0.35 [0.31, 0.39]).

## DISCUSSION

Using three-dimensional acoustic localisation and source separation techniques, we revealed how male social strategies in an arboreal reed frog unfold over several nested spatial and temporal scales. Male *H. sp. A* appear to return to the same arboreal calling sites across nights, with some locational drift over time and with 3D social structure responding flexibly to nightly variation in caller densities. Within nights, choruses formed as males descend to their calling sites at nightfall from presumed daytime refugia higher in the canopy. Once at the chorus, male call-timing strategies were flexibly adapted to varied caller densities, and key call-timing interactions were spatially structured. These results highlight that bioacoustic pipelines relying on consumer-grade hardware can allow non-invasive tracking of vocalizing individuals in space and time and accurately assign calls to different callers. Thus these approaches represent a highly cost-effective means of gaining a uniquely detailed and holistic view of the dynamics of wild animal societies, including those occupying complex nocturnal environments.

### Chorus spatial structure

In acoustically signalling animals, signal transmission, masking, and neighbour interactions are strongly distance-dependent (Naguib and Wiley, 2001). Thus, space fundamentally structures communication networks and communication strategies must include a spatial component (Reichert et al., 2021; Snijders and Naguib, 2017). Consistent with this framework, individuals occupied non-random positions, calling from a relatively constrained vertical stratum (∼3 m) while maintaining regular nearest-neighbour spacing (∼5 m). Such regular spacing is widely documented in anuran choruses and inter-male spacing is typically maintained via acoustic signals (Tárano, 2009; Wilczynski & Brenowitz, 1988; Brenowitz, 1989; Gerhardt et al., 1989). Spacing and specific site use are thought to reduce acoustic interference from nearby rivals, and secure access to high-quality calling and oviposition microhabitats, all of which can influence mating success (Wells, 1977).

Importantly, spatial structure was not fixed. Males showed calling site fidelity over shorter timescales, but this fidelity slowly degraded over time. This suggests that calling locations were repeatedly reused rather than selected randomly each night, but with some drift over time in the precise locations chosen. Furthermore, we uncovered important social drivers of changes in chorus structure across nights. As seen in other frogs, as the number of calling males increased, inter-individual distances decreased (Dyson & Passmore, 1992; Gerhardt et al., 1989). This suggests that individuals compress spacing rather than exclude additional competitors, at least to the densities observed. Thus, chorus spatial structure seems to arise as a constant negotiation among local rivals, and the terms of this negotiation adapt to changing social dynamics across nights to allow signal efficacy to be maintained under increasing density (Gerhardt et al., 1989; Wilczynski & Brenowitz, 1988). At the same time, weak evidence for upward shifts in calling height suggests that vertical displacement may provide an additional axis along which competition can be alleviated.

The magnitude and duration of this fidelity are consistent with a “structured but dynamic” model of chorus organisation described in other anurans (e.g. Tárano, 2009), and contrast with strongly territorial systems in which individuals maintain fixed, defended sites over extended periods (e.g. Ringler et al., 2009). Instead, frogs in this system appear to reuse space without strict territorial boundaries, aligning more closely with non-territorial but site-faithful systems in which individuals exhibit spatial consistency despite substantial overlap and limited aggression (Señaris et al., 2023). The gradual decay in spatial consistency over approximately one week further suggests that familiarity-based space use is flexible rather than fixed. Such patterns are consistent with models of optional resettlement, in which individuals balance the benefits of familiar space against changing environmental and social conditions (Piper, 2011). Calling locations likely vary in quality due to microhabitat structure, acoustic properties, and proximity to breeding resources, but may be difficult to evaluate prospectively, favouring repeated use of known sites with occasional repositioning. Although at this site it was not apparent that calling locations were limited on the horizontal axes due to the density of available habitat, there may have been a vertical limitation as the understory vegetation declined in density above 3 to 4 m. In future research, we suggest the addition of lidar or photogrammetry methods to map the distribution of vegetation, to directly analyse the impact of habitat availability on chorus structuring.

### Individual Movement Within Nights

Vertical movements of calling males were best described by three discrete behavioural states reflecting a gradient of spatial displacement: Stationary, characterised by minimal positional change consistent with settled calling; Local Repositioning, representing low-to-moderate fine-scale positional adjustments; and Active Displacement, reflecting large-scale vertical movement consistent with canopy transit between calling bouts. These states differed systematically in both step length magnitude and mean calling height, supporting their biological validity as distinct phases of nightly activity. The distribution of calling heights across behavioural states supports a model of active canopy descent to calling sites. Frogs in active displacement vocalised across the widest range of elevations, consistent with calls produced during transit, whereas stationary individuals converged at lower and more consistent heights, indicating the selection of specific calling positions. Local Repositioning represents an intermediate phase of fine-scale adjustment prior to settlement. When combined with the temporal progression in state composition, these patterns reveal a structured nightly sequence: frogs begin in active displacement, rapidly descending from higher positions, and subsequently transition through a period of repositioning before stabilising in calling sites.

This consistent pattern of descent suggests that individuals begin the night away from calling sites and move toward breeding habitats as chorusing activity intensifies. One plausible explanation is that frogs occupy higher arboreal refugia during the day before descending toward breeding sites near the water surface at the onset of reproductive activity. Higher arboreal positions may reduce exposure to pond-associated predators, particularly snakes, which are major predators of tropical anurans. Frog density declines steeply above the lower forest strata (0 – 3 m) both at this site and across other tropical forest systems (Basham et al., 2024; Basham & Scheffers, 2020), suggesting that even modest upward movements may substantially reduce overlap with areas of concentrated frog activity and associated predator encounter risk. This potential refuge benefit occurs despite increasingly harsh microclimatic conditions with height, including elevated temperatures and desiccation risk toward the canopy (Basham & Scheffers, 2020). Such movements may reflect a trade-off between safety and reproductive opportunity, with calling sites positioned closer to oviposition habitats but associated with elevated competition and predator exposure. The initial phase of active displacement may therefore represent a period of spatial negotiation, in which individuals assess the distribution and density of competitors before settling into calling positions, consistent with the density-dependent spatial adjustments observed at the chorus level. Critically, these results demonstrate that acoustic triangulation coupled with hidden Markov models can resolve fine-scale vertical movement and behavioural state transitions in free-ranging anurans at a spatial and temporal resolution that is otherwise extremely difficult to achieve.

### Fine-scale Call-Timing Interactions at the Chorus

*H. sp. A* exhibits novel call-timing behaviour not seen elsewhere in the frog chorusing literature. Its extremely brief standard call (mean = 0.033 s; Fig. 2a) and lengthy call period (mean = 12.8 s; Fig. 2c) combine to generate an exceptionally low duty cycle per caller, even in dense choruses (typical duty cycle = 0.003). Females typically prefer higher call rates and duty cycles (Ryan & Keddy-Hector, 1992), suggesting that such a low duty cycle may have arisen in response to countervailing ecological pressures. Avoidance of detection by eavesdropping predators and parasites seems a particularly promising candidate pressure (Ryan et al., 1982) that has selected for reduced duty cycles in various signalling taxa (Bernal & Page, 2023).

Regarding call-timing interactions, alternation in most frogs results from rapid responses (∼50 ms to a few hundred ms) to rivals’ call offsets (Larter & Ryan, 2025; Klump & Gerhardt, 1992). In 2-male choruses, this leads to lopsided alternation (often called ‘entrainment’: Grafe, 2005), in which the call of one male is followed rapidly by the other, which is then followed by an extended silence before the first male calls again. In 2-male choruses of frog species that call at very high rates, back-to-back mutual entrainment responses (response delays of ∼50 ms) can generate rapid anti-phase alternation (e.g., *Hyla arborea* and *Dryophytes cinereus*: (Gerhardt & Huber, 2002; Klump & Gerhardt, 1992)). However, *H. sp. A* is unique in maintaining near-perfect anti-phase alternation in 2-male choruses despite lengthy response latencies (∼4 s) and low duty cycles (Fig. 2b, upper). This suggests their call-timing mechanism may function quite differently from those that facilitate rapid gap-detection responses in other frogs (Larter & Ryan, 2025). Interestingly, limited data on another African Hyperoliid frog, *Kassina cassinoides*, suggests it may exhibit a similar pattern of anti-phase alternation with a relatively low duty cycle ([∼0.2 s call duration]/[∼2.5 s call period] = [duty cycle of ∼0.08]) (Grafe, 2005). Thus, Hyperoliids, or African frogs generally, may subvert generalizations based primarily on well-studied frogs from the Americas.

Finally, *H. sp. A* exhibits highly flexible call-timing responses to rivals’ calls. Callers exhibit two discrete temporal response types; rapid responses with response latencies of ∼0.16 s, and slow responses with response latencies of ∼3.06 s (Fig. 2c). Males flexibly tailor their use of different response types to the social environment, with rapid responses used more frequently in more competitive contexts such as when more rivals are calling, and when responding to nearer rivals (Fig. 5). Furthermore, latencies of slow responses decreased as more rivals called nearby. This graded latency variation within these discrete response types thus demonstrates hierarchical flexibility in call-timing responses that is socially mediated. This flexibility led to rather varied call periods, with the mean coefficient of variation in call period being 0.33. Frogs in general exhibit quite variable call periods (Larter & Ryan, 2025), but *H. sp. A* is among the more variable frog species reported.

Socially mediated flexibility in discrete call-timing responses to rivals has been reported in other frog species, but the mechanism by which it is accomplished in *H. sp. A* appears to be novel. For instance, túngara frogs can alternate with rivals’ calls or overlap them in a highly stereotyped way, and the prevalence of these responses varies across social environments (Larter et al., 2026). However, both discrete response types are driven by the same temporal response, a rapid (∼50-150 ms) response, but with this response triggered by different parts of rivals’ calls (Larter & Ryan, 2024a). Thus, túngara frogs exhibit socially mediated flexibility in *which perceptual call-triggers* are responded to. Conversely, *H. sp. A* clearly exhibits flexibility in *which discrete temporal response* is employed when responding to the same perceptual call trigger (the onset, offset, or peak amplitude of their short calls). Thus, *H. sp. A* exhibits a novel form of temporal flexibility (but see Reichert, 2012) for another potential, though ambiguous, example: *Dendropsophus ebracattus*). This contributes to our growing picture of the remarkable versatility and responsiveness of frog call-timing strategies (Larter & Ryan, 2025).

Currently the function of the two call-timing response types remains mysterious. The large and stable nearest-neighbour distances, and apparent between-night stability in calling sites in *H. sp. A* choruses, suggest a degree of spatial intolerance. Thus, rapid responses to rivals could function as an aggressive signal to maintain inter-male spacing (Brenowitz, 1989), in the same way that intentional song overlap between nearby rivals may signal aggression in birds (Naguib & Mennill, 2010). Indeed, rapid responses are employed primarily towards nearer neighbours, and in denser choruses, which is consistent with aggressive calls in other frog species (Dyson et al., 2013; Wells, 1977). However, playback experiments will be needed to uncover the functions of the different call-timing responses, and single- and multi-note calls, in *H. sp. A*.

### Conclusions

Here, we were able to simultaneously reveal several key facets of the social structure and communication ecology of a hitherto unknown African reed frog, *Hyperolius sp. A*. Importantly, the technological advance here lies not simply in localising calls, but in resolving repeated calls to individual callers in three-dimensional space and through time. This allowed individual calling histories to be reconstructed across the evening, revealing that wild social dynamics are intertwined across nested spatiotemporal scales, and generated key insights into the social strategies of this species and anurans, generally. The breadth and depth with which we were able to illuminate the social structure of this previously unknown species demonstrate the power of novel acoustic technologies for studying previously inaccessible acoustically signalling systems in the wild, with relevance for many other taxa such as insects, birds, bats, primates, and cetaceans.

A major challenge for extending these approaches is the trade-off between the resolution of behavioural information and the volume of acoustic data that can be processed. The beamforming and clustering approach used here allowed us to reconstruct highly resolved individual calling histories, but was labour intensive. Conversely, convolutional neural networks and other automated detection methods can process much larger acoustic datasets, but may fail to detect or correctly classify low-quality, overlapping or otherwise atypical calls, potentially limiting their application where complete call histories are required. An important technological direction will therefore be the integration of scalable automated call detection with high-resolution acoustic localisation, retaining the individual-level spatial and temporal information demonstrated here while substantially increasing the volume of data that can be analysed.

## REFERENCES

Alexander, R. D. (1974). The Evolution of Social Behavior. Annual Review of Ecology and Systematics, 5(1), 325–383. 10.1146/annurev.es.05.110174.001545

Basham, E. W., Baecher, J. A., Klinges, D. H., & Scheffers, B. R. (2022). Vertical stratification patterns of tropical forest vertebrates: A meta-analysis. Biological Reviews, 98(1), 99–114. 10.1111/brv.12896

Basham, E. W., & Scheffers, B. R. (2020). Vertical stratification collapses under seasonal shifts in climate. Journal of Biogeography, 47(9), 1888–1898. 10.1111/jbi.13857

Basham, E. W., Scheffers, B. R., Nakamura, A., Bamba-Kaya, A., & Jongsma, G. F. M. (2024). Vertical niche and trait associations in Central African amphibians. Biotropica, 56(4), e13349. 10.1111/btp.13349

Bates, D., Mächler, M., Bolker, B. M., & Walker, S. C. (2015). Fitting linear mixed-effects models using lme4. Journal of Statistical Software, 67(1), 1–48. 10.18637/jss.v067.i01

Bernal, X. E., & Page, R. A. (2023). Tactics of evasion: Strategies used by signallers to deter eavesdropping enemies from exploiting communication systems. Biological Reviews, 98(1), 222–242. 10.1111/brv.12904

Brenowitz, E. A. (1989). Neighbor Call Amplitude Influences Aggressive Behavior and Intermale Spacing in Choruses of the Pacific Treefrog (*Hyla regilla*). Ethology, 83(1), 69–79. 10.1111/j.1439-0310.1989.tb00520.x

Brush, J. S., & Narins, P. M. (1989). Chorus dynamics of a neotropical amphibian assemblage: Comparison of computer simulation and natural behaviour. Animal Behaviour, 37, 33–44. 10.1016/0003-3472(89)90004-3

Calsbeek, R., Zamora-Camacho, F. J., & Symes, L. B. (2022). Individual contributions to group chorus dynamics influence access to mating opportunities in wood frogs. Ecology Letters, 25(6), 1401–1409. 10.1111/ele.14002

Dougherty, R. P., Pulica, R. M., & Caldwell, M. S. (2022). Multi-night territorial behavior, chorus attendance, and mating success in red-eyed treefrogs. Ethology, 128, 608–619. 10.1111/eth.13321

Dyson, M. L., Reichert, M. S., & Halliday, T. R. (2013). Contests in amphibians. In I. Hardy & M. Briffa (Eds.), Animal Contests (pp. 228–257). Cambridge University Press.

Dyson, M. L., & Passmore, N. I. (1992). Inter-male Spacing and Aggression in African Painted Reed Frogs, *Hyperolius marmoratus*. Ethology, 91(3), 237–247. 10.1111/j.1439-0310.1992.tb00865.x

Friedl, T. W. P., & Klump, G. M. (2005). Sexual selection in the lek-breeding European treefrog: Body size, chorus attendance, random mating and good genes. Animal Behaviour, 70(5), 1141–1154. 10.1016/j.anbehav.2005.01.017

Gerhardt, H. C., & Huber, F. (2002). Acoustic communication in insects and anurans: Common problems and diverse solutions. University of Chicago Press.

Gerhardt, H. C., Diekamp, B., & Ptacek, M. (1989). Inter-male spacing in choruses of the spring peeper, Pseudacris (Hyla) crucifer. Animal Behaviour, 38(6), 1012–1024. 10.1016/S0003-3472(89)80140-X

Grafe, T. U. (2005). Anuran choruses as communication networks. In P. K. McGregor (Ed.), Animal Communication Networks (pp. 277–300). Cambridge University Press.

Greenfield, M. D. (1994). Cooperation and conflict in the evolution of social interactions. Annual Review of Ecology and Systematics, 25(1), 97–126. 10.1146/annurev.es.25.110194.000525

Hödl, W. (1977). Call differences and calling site segregation in anuran species from central Amazonian floating meadows. Oecologia, 28(4), 351–363. 10.1007/BF00345990

Jones, D. L., Jones, R. L., & Ratnam, R. (2014). Calling dynamics and call synchronization in a local group of unison bout callers. Journal of Comparative Physiology A: Neuroethology, Sensory, Neural, and Behavioral Physiology, 200(1), 93–107. 10.1007/s00359-013-0867-x

Jones, D. L., & Ratnam, R. (2009). Blind location and separation of callers in a natural chorus using a microphone array. The Journal of the Acoustical Society of America, 126(2), 895–910. 10.1121/1.3158924

Klump, G. M., & Gerhardt, H. C. (1992). Mechanisms and function of call-timing in male-male interactions in frogs. In Playback and Studies of Animal Communication (pp. 153–174). Springer.

Krebs, J. R., & Davies, N. B. (1978). Behavioural Ecology: An Evolutionary Approach. Blackwell Publishing.

Kuhn, H. W. (1955). The Hungarian method for the assignment problem. Naval Research Logistics Quarterly, 2(1–2), 83–97. 10.1002/nav.3800020109

Kuznetsova, A., Brockhoff, P. B., & Christensen, R. H. B. (2017). lmerTest Package: Tests in Linear Mixed Effects Models. Journal of Statistical Software, 82(13), 1–26. 10.18637/jss.v082.i13

Larter, L. C., Cushing, C. W., & Ryan, M. J. (2026). Cadences of the collective: Conspecific stimulation patterns interact with endogenous rhythms to cue socially mediated response shifts. Journal of Experimental Biology, 229(1), jeb250982. 10.1242/jeb.250982

Larter, L. C., & Ryan, M. J. (2024a). Sensory-motor tuning allows generic features of conspecific acoustic scenes to guide rapid, adaptive, call-timing responses in túngara frogs. Proceedings of the Royal Society B: Biological Sciences, 291(2031), 20240992. 10.1098/rspb.2024.0992

Larter, L. C., & Ryan, M. J. (2024b). Túngara frog call-timing decisions arise as internal rhythms interact with fluctuating chorus noise. Behavioral Ecology, 35(4), arae034. 10.1093/beheco/arae034

Larter, L. C., & Ryan, M. J. (2025). The Variability and Malleability of Frog Call-Timing Mechanisms are Neglected in Traditional Call-Timing Models. *Integrative and Comparative Biology*, icaf041. 10.1093/icb/icaf041

Michelot, T., Langrock, R., & Patterson, T. A. (2016). moveHMM: An R package for the statistical modelling of animal movement data using hidden Markov models. Methods in Ecology and Evolution, 7(11), 1308–1315. 10.1111/2041-210X.12578

Morse, D. H. (1980). Behavioral mechanisms in ecology. Harvard University Press.

Naguib, M., & Mennill, D. J. (2010). The signal value of birdsong: Empirical evidence suggests song overlapping is a signal. Animal Behaviour, 80(3), e11–e15. 10.1016/j.anbehav.2010.06.001

Naguib, M., & Wiley, R. H. (2001). Estimating the distance to a source of sound: Mechanisms and adaptations for long-range communication. Animal Behaviour, 62(5), 825–837. 10.1006/anbe.2001.1860

Piper, W. H. (2011). Making habitat selection more “familiar”: A review. Behavioral Ecology and Sociobiology, 65(7), 1329–1351. 10.1007/s00265-011-1195-1

Pohle, J., Langrock, R., Van Beest, F. M., & Schmidt, N. M. (2017). Selecting the Number of States in Hidden Markov Models: Pragmatic Solutions Illustrated Using Animal Movement. *Journal of Agricultural*, Biological and Environmental Statistics, 22(3), 270–293. 10.1007/s13253-017-0283-8

R Core Team. (2023). R: A language and environment for statistical computing. R Foundation for Statistical Computing.

Reichert, M. S. (2012). Call timing is determined by response call type, but not by stimulus properties, in the treefrog *Dendropsophus ebraccatus*. Behavioral Ecology and Sociobiology, 66(3), 433–444. 10.1007/s00265-011-1289-9

Reichert, M. S., Enriquez, M. S., & Carlson, N. V. (2021). New Dimensions for Animal Communication Networks: Space and Time. Integrative and Comparative Biology, 61(3), 814–824. 10.1093/icb/icab013

Rhinehart T. A., Chronister L. M., Devlin T., & Kitzes J. (2020) Acoustic localization of terrestrial wildlife: Current practices and future opportunities. Ecology and Evolution, 10: 6794–6818. 10.1002/ece3.6216

Richards, D. G., & Wiley, R. H. (1980). Reverberations and Amplitude Fluctuations in the Propagation of Sound in a Forest: Implications for Animal Communication. The American Naturalist, 115(3), 381–399.

Ringler, M., Ursprung, E., & Hödl, W. (2009). Site fidelity and patterns of short- and long-term movement in the brilliant-thighed poison frog *Allobates femoralis* (Aromobatidae). Behavioral Ecology and Sociobiology, 63(9), 1281–1293. 10.1007/s00265-009-0793-7

Ryan, M. J., & Keddy-Hector, A. (1992). Directional Patterns of Female Mate Choice and the Role of Sensory Biases. The American Naturalist, 139, S4–S35.

Ryan, M. J., Tuttle, M. D., & Rand, A. S. (1982). Bat Predation and Sexual Advertisement in a Neotropical Anuran. The American Naturalist, 119(1), 136–139.

Schwartz, J. J. (1993). Male calling behavior, female discrimination and acoustic interference in the Neotropical treefrog Hyla microcephala under realistic acoustic conditions. Behavioral Ecology and Sociobiology, 32(6), 401–414. 10.1007/BF00168824

Señaris, C., Lampo, M., Rodríguez-Contreras, A., & Velásquez, G. (2023). Breeding site fidelity of the critically endangered toad *Atelopus cruciger* (Anura: Bufonidae): Implications for its conservation. Salamandra, 59(3), 217–228.

Sievert, C., Parmer, C., Hocking, T., Chamberlain, S., Ram, K., Corvellec, M., & Despouy, P. (2015). plotly: Create Interactive Web Graphics via “plotly.js” (p. 4.12.0) [Dataset]. 10.32614/CRAN.package.plotly

Snijders, L., & Naguib, M. (2017). Communication in Animal Social Networks. In Advances in the Study of Behavior (Vol. 49, pp. 297–359). Elsevier. 10.1016/bs.asb.2017.02.004

Tárano, Z. (2009). Structure of Transient Vocal Assemblages of *Physalaemus fischeri* (Anura, Leiuperidae): Calling Site Fidelity and Spatial Distribution of Males. South American Journal of Herpetology, 4(1), 43. 10.2994/057.004.0105

Webber, Q. M. R., Albery, G. F., Farine, D. R., Pinter-Wollman, N., Sharma, N., Spiegel, O., Vander Wal, E., & Manlove, K. (2023). Behavioural ecology at the spatial–social interface. Biological Reviews, 98(3), 868–886. 10.1111/brv.12934

Wells, K. D. (1977). The social behaviour of anuran amphibians. Animal Behaviour, 25, 666–693. 10.1016/0003-3472(77)90118-X

Wells, K. D. (1978). Territoriality in the green frog (Rana clamitans): Vocalizations and agonistic behaviour. Animal Behaviour, 26, 1051–1063. 10.1016/0003-3472(78)90094-5

Wilczynski, W., & Brenowitz, E. A. (1988). Acoustic cues mediate inter-male spacing in a neotropical frog. Animal Behaviour, 36(4), 1054–1063. 10.1016/S0003-3472(88)80065-4

