## supplemental for "4-Dimensional Chess: Acoustic Localisation Reveals Nested Spatio-temporal Strategies in an Arboreal Communication Network"

**Supplementary Materials**

**Supplemental Methods 1**: Null model of inter-night spatial consistency

To generate an empirical null expectation for inter-night spatial consistency, all triangulated frog locations across all nights (*n* = 7,842) were pooled to represent the observed calling space. Locations were discretised into a 0.5-m voxel grid, and duplicate occurrences within the same voxel were removed, resulting in 1,175 unique occupied voxels. The resulting voxel pool therefore represented locations within the study area at which calling frogs had been observed, while preventing frequently occupied locations from receiving greater weight simply because they contained more call detections.

Null locations were randomly sampled from these occupied voxels, with each voxel having equal probability of selection. For each inter-night comparison, random centroid sets were generated while retaining the observed numbers of callers, and spatial consistency was calculated using the same optimal one-to-one matching procedure applied to the observed nightly centroid sets. When the numbers of callers differed between nights, matching was restricted to the maximum possible number of pairs and unmatched locations were excluded. Repeated randomisations generated the null distribution of mean matched distances against which the observed inter-night spatial consistency was evaluated.

This approach tests whether frogs reused spatial locations across nights more consistently than expected from random reuse of the empirically occupied calling space, rather than assuming that all three-dimensional space within the acoustic array represented equally available calling habitat.

Hornik, K. (2004). *clue: Cluster Ensembles* (p. 0.3-68) [Dataset]. https://doi.org/10.32614/CRAN.package.clue

**Supplementary Methods 2**: Additional decision-based parameters for HMM fitting and analysis filtering.

(i) HMM initial parameter values. Initial step length parameters were selected to reflect the expected biological gradient across states: near-zero values for Stationary ($\mu$ = 0.03 m, $\sigma$ = 0.02 m), intermediate values for Local Repositioning ($\mu$ = 0.20 m, $\sigma$ = 0.15 m), and large values for Active Displacement ($\mu$ = 0.60 m, $\sigma$ = 0.50 m; Supplementary Table S2). The two-state model was fitted solely for AIC comparison and is not interpreted biologically.

(ii) Transition probability matrix. Transition probabilities were derived from the multinomial logit parameterisation implemented in moveHMM, with no constraints placed on the transition probability matrix during fitting. Log-odds regression coefficients are reported in Supplementary Table S4; back-transformed transition probabilities are provided in the same table.

(iii) State decoding. Following model fitting, posterior state probabilities were extracted using the stateProbs() function in moveHMM. The most probable state at each time step was assigned via which.max() applied across the posterior probability matrix, equivalent to Viterbi decoding under the fitted model.

(iv) Pseudo-residual diagnostics. Model fit was assessed via pseudo-residuals computed using the plotPR() function in moveHMM. Pseudo-residuals were inspected for departures from normality and temporal autocorrelation as indicators of model misspecification.

(v) Filter justification. Two sequential filters were applied prior to statistical modelling. The threshold of five observations per frog-night was chosen as the minimum number of fixes required for meaningful behavioural state sequence inference. The 1,800 s (30-minute) minimum session duration threshold was selected to ensure sufficient temporal coverage to detect within-night state transitions

**Supplemental Methods 3**: K-Means Clustering to Optimally Separate Rapid and Slow Call-Timing Responses

The distribution of log-transformed response delays showed clear bimodality (see Figure 4, in-text). To mathematically determine the optimal threshold separating the two modes evident in this distribution, we implemented k-means clustering with the ‘kmeans()’ function from the *stats* R package. We clustered the entire distribution of log-transformed response delays (n = 2, 158), looking for 2 centers, and trying 50 random starting points. This gave us two centers that closely matched our subjective assessment of the peaks in the distribution. We then found the middle point between these centers by taking their mean, and used that as our threshold. We back-transformed these centers and thresholds to the original linear scale, giving us two centers in the distribution at 0.16s and 3.06s, separated by a threshold at 0.69s. Visually, this separated the two peaks well (see Figure 4, in-text). We thus defined responses with onset delays less than 0.69s as rapid responses, and those with onset delays greater than or equal to 0.69 as slow responses. The code for these operations can be found in the submitted R files.

### Supplementary Tables

**Supplementary Table 1:** AIC-based model selection comparing two- and three-state Hidden Markov Models fitted to vertical step-length data from *Hyperolius sp. A*.

| Model | Number of states | AIC | ΔAIC | Selected |
|---|---|---|---|---|
| Two-state HMM | 2 | -427.17 | 27.96 | No |
| Three-state HMM | 3 | -455.13 | 0 | Yes |

Note: ΔAIC > 10 is considered decisive evidence against the higher-AIC model. The three-state model was retained on both statistical and biological grounds (Pohle et al., 2017).

**<u>Supplementary Table 2:</u>** Initial parameter values used for HMM fitting. State means (μ) and standard deviations (σ) for the gamma step-length distribution.

| State | Label | μ (m) | σ (m) |
|---|---|---|---|
| 1 | Stationary | 0.03 | 0.02 |
| 2 | Slow Movement | 0.2 | 0.15 |
| 3 | Active Displacement | 0.6 | 0.5 |

Note: Initial values were chosen to reflect a plausible gradient from near-stationary to actively displaced calling behaviour.

**<u>Supplementary Table 3:</u>** Mean calling height (m) by HMM-derived behavioural state across all individuals.

| State | N fixes | Mean height (m) | SD (m) |
|---|---|---|---|
| Stationary | 279 | 3.43 | 1.2 |
| Slow Repositioning | 632 | 2.98 | 1.33 |
| Active Displacement | 214 | 3.55 | 1.68 |

**Supplementary Table 4:** HMM transition probability matrix and initial state distribution. Values represent the probability of transitioning from state *i* (row) to state *j* (column) at consecutive vocalisation fixes.

| From \ To | Stationary | Slow Repositioning | Active Displacement |
|---|---|---|---|
| Stationary | 80.70% | 16.60% | 2.70% |
| Slow Repositioning | 10.00% | 87.60% | 2.30% |
| Active Displacement | 8.60% | 36.20% | 55.20% |

**Supplementary Table 5:** Estimated marginal mean effects [95% CIs] of predictor variables across our various response models, shown for a representative range of predictor values. Grayed-out cells represent predictors that did not significantly influence outcomes.

| **Predictor Variable:** | ***Model Name:*** | | | |
|---|---|---|---|---|
| | *Response_Type_GLMM* | *Rapid_Response_Latency_LMM* | *Slow_Response_Latency_LMM* | *Call_Period_LMM* |
| | **Response Variable:** | | | |
| | Prob(Rapid Response) | Rapid Response Latency (s) | Slow Response Latency (s) | Call Period (s) |
| Wait Time | **5s:** 0.12 [0.09, 0.15] | **5s:** 0.24 [0.21, 0.26] | **5s:** 3.17 [2.84, 3.54] | -- |
| | **12.5s:** 0.22 [0.18, 0.26] | **12.5s:** 0.2 [0.19, 0.22] | **12.5s:** 2.81 [2.52, 3.14] | -- |
| | **20s:** 0.37 [0.30, 0.44] | **20s:** 0.17 [0.14, 0.2] | **20s:** 2.49 [2.22, 2.8] | -- |
| Number Active Callers | **2:** 0.08 [0.05, 0.11] | **2:** 0.19 [0.15, 0.22] | **2:** 4.19 [3.67, 4.77] | **2:** 13.26 [12.29, 14.22] |
| | **4:** 0.16 [0.13, 0.19] | **4:** 0.2 [0.18, 0.22] | **4:** 3.08 [2.77, 3.44] | **4:** 12.85 [12.09, 13.6] |
| | **6:** 0.29 [0.24, 0.35] | **6:** 0.21 [0.19, 0.23] | **6:** 2.27 [2.0, 2.58] | **6:** 12.44 [11.55, 13.33] |
| Inter-Caller Distance | **5m:** 0.43 [0.34, 0.53] | **5m:** 0.19 [0.16, 0.21] | **5m:** 3.46 [3.07, 3.9] | **5m:** 12.41 [11.58, 13.23] |
| | **12.5m:** 0.16 [0.13, 0.19] | **12.5m:** 0.21 [0.2, 0.23] | **12.5m:** 2.9 [2.6, 3.23] | **12.5m:** 12.8 [12.04, 13.56] |
| | **20m:** 0.04 [0.03, 0.07] | **20m:** 0.24 [0.2, 0.27] | **20m:** 2.43 [2.13, 2.77] | **20m:** 13.19 [12.29, 14.1] |
| Response Type | -- | -- | -- | **Rapid**: 12.27s [11.44, 13.1] |
| | -- | -- | -- | **Slow**: 13.22 [12.49, 13.99] |

### Supplementary Figures

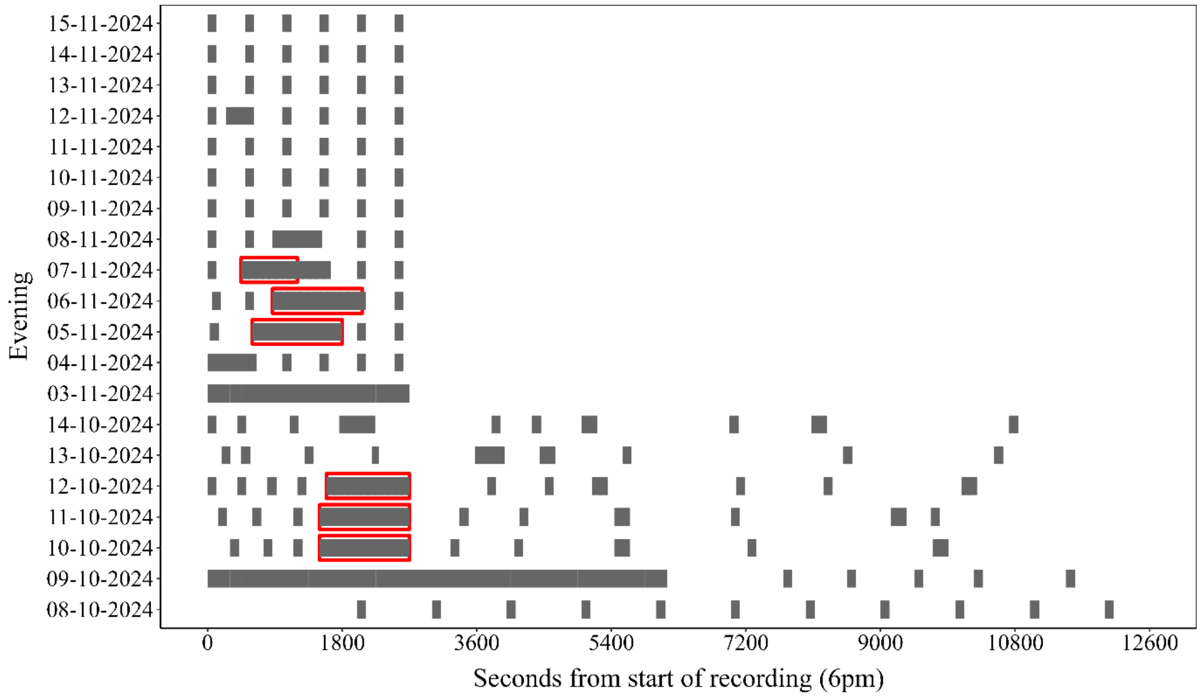


**Figure S1:** Gantt chart showing the distribution of audio recording windows where calls were triangulated. Session 3 recordings (08/10/2024 - 14/10/2024 were 3.5 hrs in length (12600 seconds), while Session 4 recordings (03/11/2024 - 15/11/2024 were 45 minutes in length (2700 seconds). Audio sections used for interaction analyses are highlighted in red.

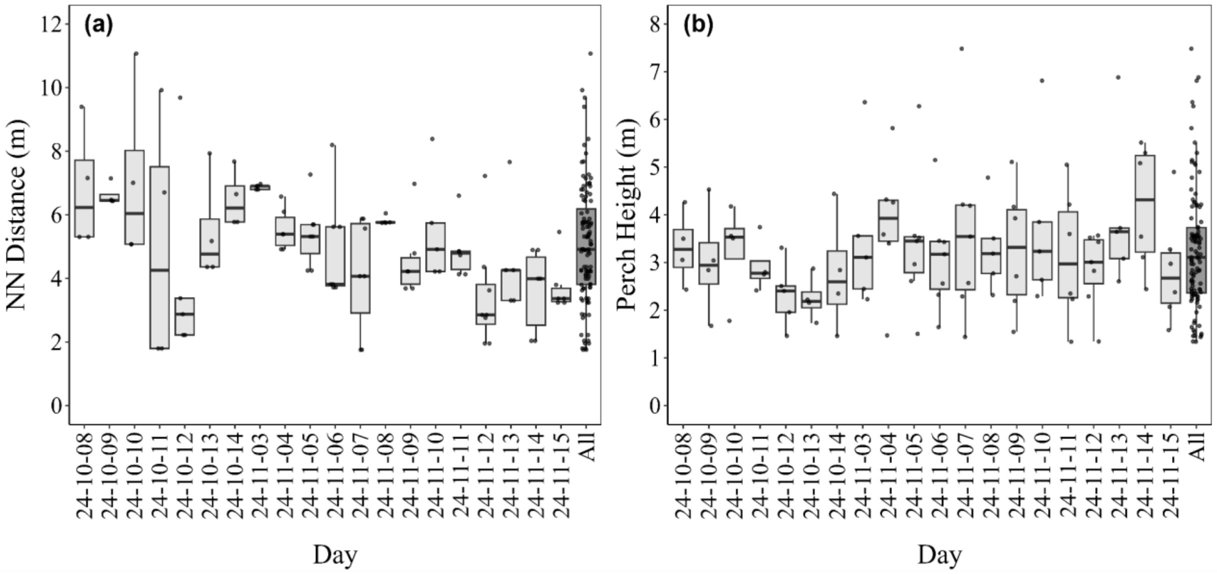


**Figure S2:** Spatial structure of the *H. sp. A* calling chorus across 20 recording nights (Session 3 (October) and session 4 (November). (a) Nearest-neighbour distance between simultaneously calling individuals for each frog-night. (b) Calling height above ground for each frog-night. One observation at 10.2 m is excluded from the plot axis range for clarity. Boxes show median and interquartile range; individual points are jittered for clarity; the "All" category pools all frog-nights across both sessions.

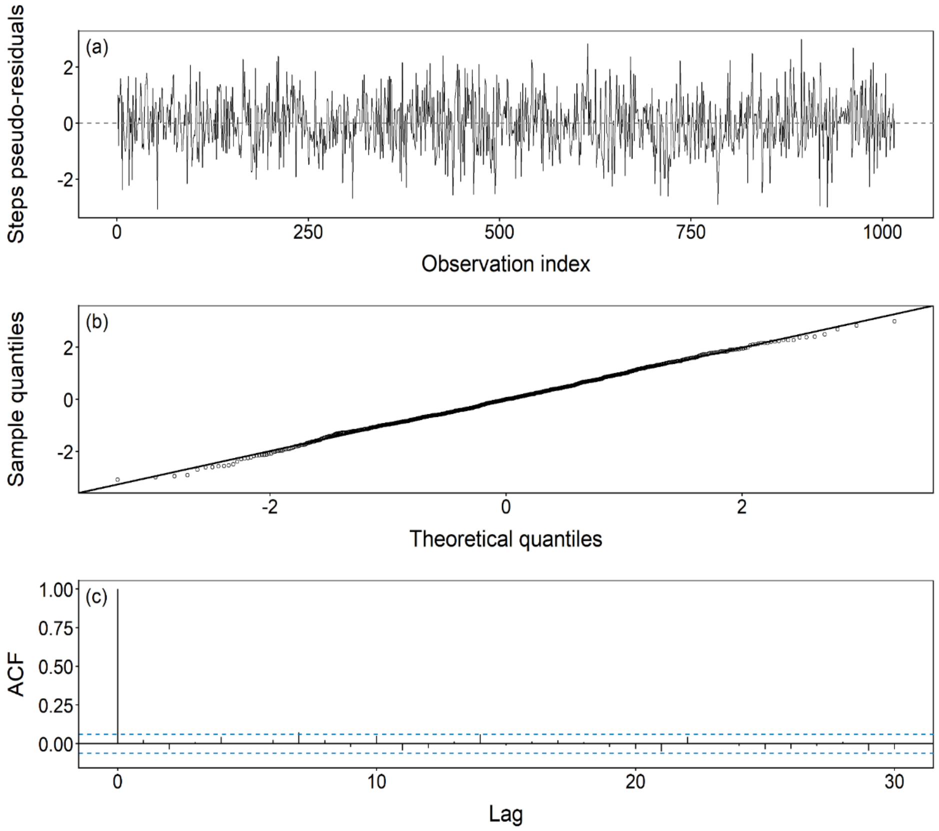


**Figure S3.** Pseudo-residual diagnostics for the three-state HMM: pseudo-residuals plotted against observation index, quantile-quantile plot of pseudo-residuals against theoretical normal quantiles, and, autocorrelation function (ACF) of pseudo-residuals. Approximate normality and low autocorrelation indicate adequate model fit.

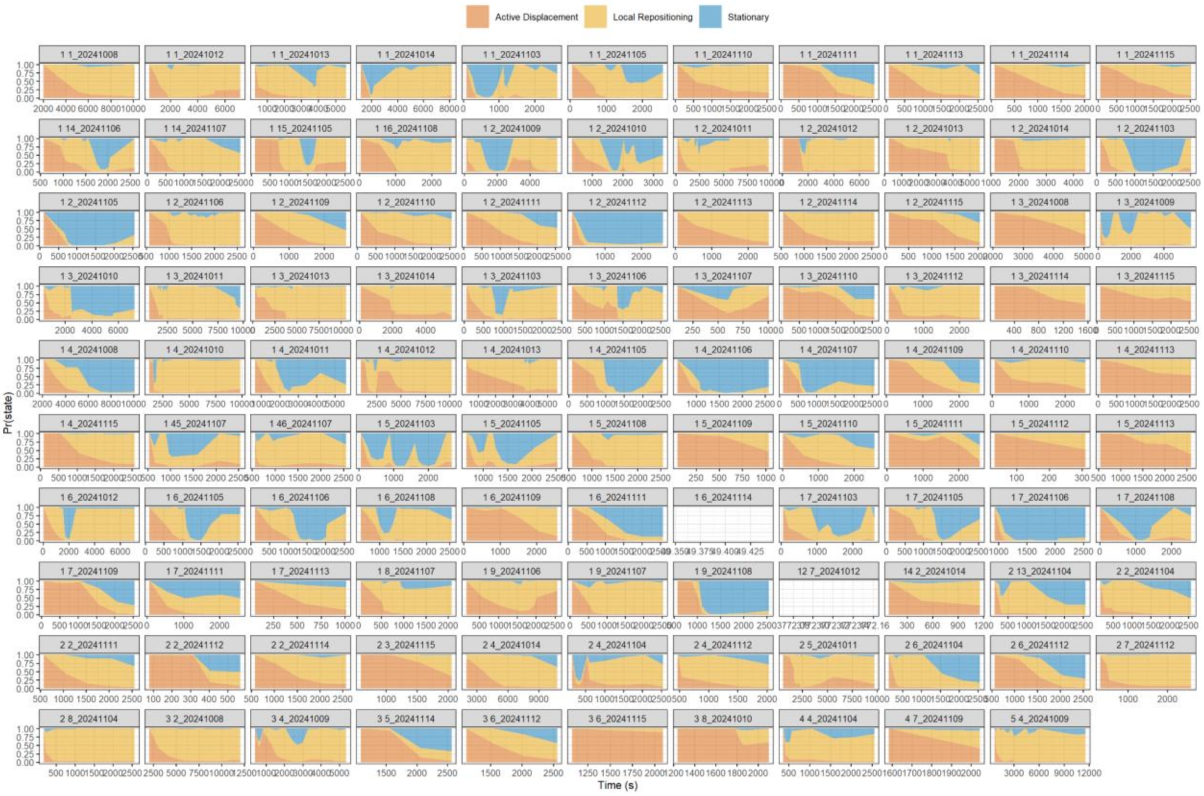


**Figure S4.** Posterior state probabilities over time for all 109 frog-nights. Coloured ribbons represent the probability of each behavioural state at each time step as estimated by the three-state HMM: Active Displacement (red), Local Repositioning (orange), and Stationary (blue). Each panel represents one frog-night (ID = individual × date).

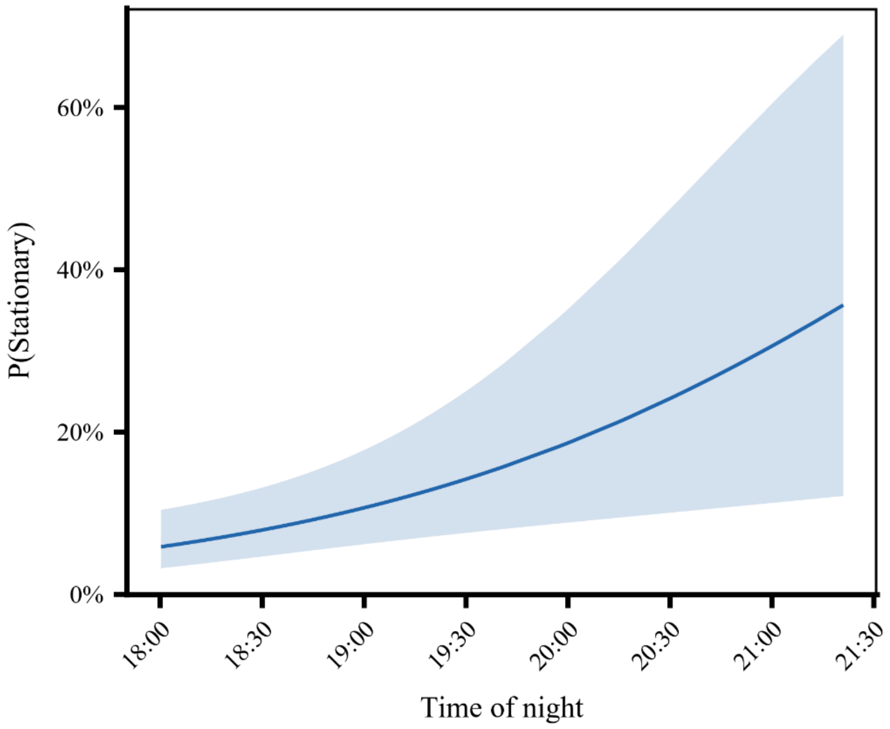


**Figure S5.** Predicted probability of Stationary, rising from ~6% at session onset to ~35% by 3.5 hours. All predictions are marginal fixed-effect estimates; shaded regions represent 95% confidence intervals.
